# Changes in prefrontal GABA and glutamate through adolescence supports excitation/inhibition balance

**DOI:** 10.1101/2022.04.25.489387

**Authors:** Maria I. Perica, Finnegan J. Calabro, Bart Larsen, Will Foran, Victor E. Yushmanov, Hoby Hetherington, Brenden Tervo-Clemmens, Chan-Hong Moon, Beatriz Luna

**Affiliations:** Department of Psychology, University of Pittsburgh; Department of Psychiatry, University of Pittsburgh; Department of Biomedical Engineering, University of Pittsburgh; Department of Psychiatry, Perelman School of Medicine, University of Pennsylvania; Department of Radiology, University of Pittsburgh; Resonance Research Incorporated, Billerica MA; Department of Radiology, University of Missouri, Columbia MO; Department of Psychiatry, Massachusetts General Hospital/Harvard Medical School

## Abstract

Animal and human postmortem studies provide evidence for changes in gamma-aminobutyric acid (GABA) and glutamate in prefrontal cortex (PFC) during adolescence, suggesting shifts in excitation and inhibition balance consistent with critical period plasticity. However, how GABA and glutamate change through adolescence and how the balance of these inhibitory and excitatory neurotransmitters changes is not well understood *in vivo* in humans. High field (7 Tesla) Magnetic Resonance Spectroscopic Imaging was used to investigate age-related changes in the balance of GABA/creatine (Cr) and glutamate/Cr in multiple developmentally-relevant regions of PFC in 144 10 to 30-year-olds. Results indicated a homogenous pattern of age-related Glu/Cr decreases across PFC regions, while age-related changes in GABA/Cr were heterogenous, with a mix of stable and decreasing age effects. Importantly, balance between glutamate/Cr and GABA/Cr in areas of prefrontal cortex increased through adolescence, suggesting the presence of critical period plasticity in PFC at this significant time of development when adult trajectories are established.

## Introduction

Adolescence is a period of significant maturation in prefrontal cortex (PFC), supporting development of cognitive control that stabilizes in adulthood (Luna et al., 2015; Sydnor et al., 2021). Synaptic pruning (Petanjek et al., 2011) and gray matter thinning in PFC continues through adolescence (Gogtay et al., 2004), while its functional (Calabro et al., 2019; Gabard-Durnam et al., 2014; Jalbrzikowski et al., 2017; Parr et al., 2021), and structural (Jalbrzikowski et al., 2017; Lebel & Beaulieu, 2011; Simmonds et al., 2014) connectivity continues to specialize supporting cognitive development (Liston et al., 2006). Underlying this transitional developmental period might be heightened neural plasticity consistent with a critical period. At the molecular level, animal and postmortem human studies show changes in GABA and glutamate in PFC (Caballero et al., 2021), which has been proposed to be consistent with a model of adolescence as a developmental critical period for executive function (Larsen & Luna, 2018).

Studies in animal model sensory systems have demonstrated that a key mechanism regulating the opening of a critical period is a shift in the excitation/inhibition (E/I) balance. In visual cortex, the critical period occurs in the first years of life, driven by maturation of inhibitory circuitry, particularly Parvalbumin (PV) neurons, which leads to the establishment of balanced excitation and inhibition, thus closing the critical period (Dorrn et al., 2010; Hensch & Fagiolini, 2005; Toyoizumi et al., 2013). More recently, it has been proposed that critical periods progress hierarchically through development, with sensory area critical periods occurring early in development, and critical periods of higher-order association cortices occurring later in development (Larsen & Luna, 2018; Reh et al., 2020). Animal and human postmortem evidence of increases in aspects of prefrontal GABAergic PV neurons (Caballero et al., 2014; Erickson & Lewis, 2002) and reductions in processes of prefrontal glutamatergic signaling (Harris et al., 2009; Henson et al., 2008; Hoftman et al., 2017) provide support for this proposal. Taken together, this literature suggests that dynamic shifts in E/I balance may be occurring during adolescence in cognitively-relevant association cortex.

To date there is still limited understanding of E/I development *in vivo* in human adolescent prefrontal cortex. Recently, a study applying pharmacological fMRI with a GABAergic benzodiazepine challenge to model changes in E/I balance through adolescence found a reduction in excitation relative to inhibition in association cortex during adolescence (Larsen et al., 2021), providing initial indirect evidence of greater inhibition through adolescence in association cortex. Initial findings from Magnetic Resonance Spectroscopy studies using 1.5T, 3T, or 4T scanners in single voxels provide some support, although findings have been inconsistent with some studies showing no age-related changes in glutamate (Blüml et al., 2013; Ghisleni et al., 2015; Gleich et al., 2015; Silveri et al., 2013) while others find age-related decreases (Shimizu et al., 2017), as well as some studies showing greater GABA levels in adults relative to adolescents (Ghisleni et al., 2015; Silveri et al., 2013).

However, these MRS studies were performed at lower field strengths, which provides some challenges to interpreting the data. First, it is more difficult to separate the overlapping signals of glutamate and its metabolic precursor, glutamine, thus undermining the ability to assess developmental change specific to glutamate. Further, GABA measurements can be confounded by overlapping macromolecule signals, which can impact measurement of GABA levels (Bell et al., 2021). Moreover, these studies typically measure changes within one large voxel (e.g., with typical voxel sizes ranging from 6 cm^3^ (Shimizu et al., 2017) to 30 cm3 (Ghisleni et al., 2015)), often leading to a reduction in the amount of gray matter relative to white matter in the voxel, is likely to influence measurement (Choi et al., 2006). Thus, while these initial results suggest that there may be important developmental changes in GABA and glutamate levels, these methodological limitations, in addition to small sample sizes (ranging from 28-56 participants), limit the ability to draw conclusions. The advent of 7T Magnetic Resonance Spectroscopic Imaging (MRSI) allows for better separation of confounding signals by doubling signal strength and improving signal-to-noise ratio (Pradhan et al., 2015), thus providing a more precise assessment of GABA and glutamate.

Here we apply 7T MRSI in a sample of 144 10-30 year-olds to assess age-related changes in GABA/Cr and glutamate/Cr. In this study, we acquired a multivoxel slice with voxel sizes of 0.9×0.9×1.0 cm, typically at least an order of magnitude smaller than previous developmental MRS studies to provide greater anatomical specificity within regions of PFC gray matter. We also assessed age-related changes in inter-subject variability, given evidence for developmental decreases in behavioral and brain functional variability (Klein et al., 2011; Montez et al., 2017; Ordaz et al., 2013; Tamnes et al., 2012). Following evidence from sensory system critical periods, we hypothesized that there would be decreases in glutamate/Cr and increases in GABA/Cr in PFC through adolescence to adulthood, resulting in a decrease in glutamate/GABA ratio and greater correlation or balance between glutamate and GABA, as well as decreases in inter-subject variability with age. While we found widespread age-related decreases in glutamate/Cr across both lateral and medial regions of the prefrontal cortex, GABA/Cr showed a regionally heterogeneous pattern of age effects, including no age-related change in some regions but decreases in others. Importantly, we also identify regions showing age-related increases in the correlation of glutamate and GABA, which might reflect greater E/I balance in adulthood as compared to adolescence. Together, the current study provides evidence for changes in the balance of glutamate and GABA through adolescence that might reflect critical period plasticity mechanisms.

## Results

### Age-related changes in metabolite levels

We characterized age-related changes in GABA/Cr and glutamate/Cr (Glu/Cr) including sex and fraction gray matter (frac GM) in the voxel as covariates in regions selected from across an MRSI slice, including dorsolateral prefrontal cortex (DLPFC), anterior cingulate cortex (ACC), anterior insula (AIns), and medial prefrontal cortex (MPFC). In regions that had both a left and a right hemisphere, data from both hemispheres was included in statistical models with hemisphere included as a covariate, and age by hemisphere interactions considered. Results showed significant age-related decreases in Glu/Cr in the DLPFC (Standardized coefficient *β* [see methods] = 0.15, Bonferroni-corrected *p* = 0.016), ACC (*β* = 0.21, *p* =0.0056), and AIns (*β* = .29, *p* = 6.5×10^−9^), and no change in the MPFC (*β* = 0.12, *p* = 0.24) (Figure 1). These relationships remained significant upon inclusion of all covariates (Supplementary Table 1). There was a main effect of hemisphere in the DLPFC (*β* = -0.40, *p* = 7.84 × 10^−6^) as well as the AIns (*β* = -0.76, *p* =2.28×10^−28^), with the left hemisphere having overall higher levels of Glu/Cr than the right. There were no significant interactions of age with hemisphere or sex in any of the regions.

**Figure 1:**
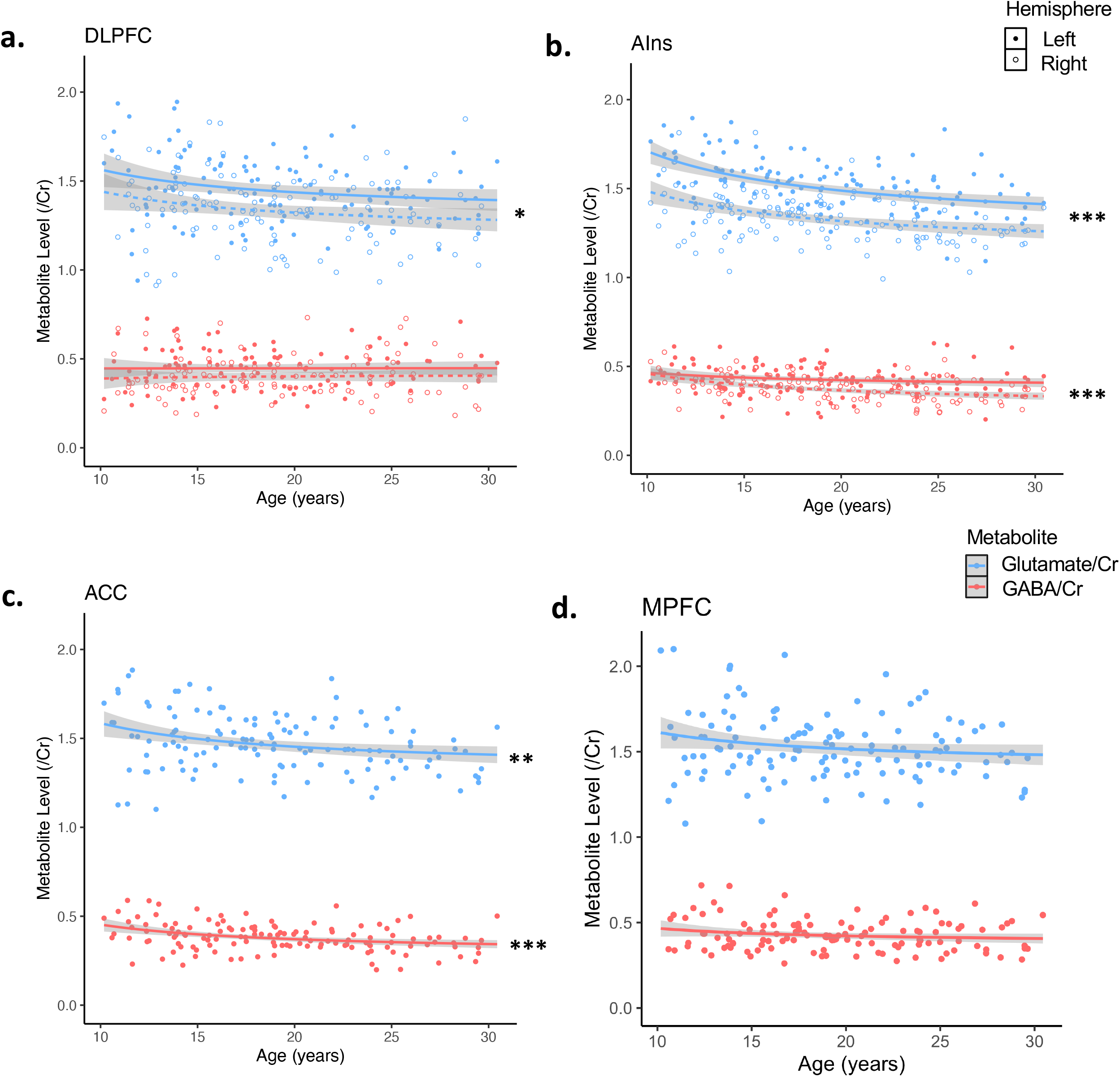
Glutamate/Cr (blue) and GABA/Cr (red) age effects across PFC regions of interest. Statistical values reflect linear mixed effects model fits for regions with both a left and right hemisphere and linear model fits for midline regions. Solid line and dots reflect left hemisphere and dashed line and open dots reflect right hemisphere where applicable. All are modeled with an inverse function. (A) left and right dorsolateral prefrontal cortex (B) left and right anterior insula (C) midline anterior cingulate cortex (d) midline medial prefrontal cortex. \**p* < .05, ** *p* < .01, *** *p* < .001. Asterisks reflect Bonferroni-corrected p-values.

Results showed significant age-related decreases in GABA/Cr in the AIns (*β* = 0.20, *p* = 9.04 ×10^−5^) and the ACC (*β* = 0.26, *p* =3.79 × 10^−4^), and no change in the MPFC (*β* = 0.11, *p* = 0.36) or DLPFC (*β* = -0.01, *p* = 1.00) (Figure 1). As with Glu/Cr, we observed a main effect of hemisphere in the DLPFC (*β* = -0.26, *p* = 0.004) and AIns (*β* = -0.49, *p* = 6.56 ×10^−8^), with higher GABA/Cr levels in the left as compared to the right hemisphere. Again, these relationships remained significant upon inclusion of all covariates (Supplementary Table 2). In the AIns, there was an uncorrected-significant age-by-hemisphere interaction, but this did not survive multiple comparisons correction (*β* = 0.16, Bonferroni-corrected *p* = 0.19). No other age-by-covariate interactions were observed.

Age-related changes in the ratio of Glu/Cr to GABA/Cr were examined for each region of interest. Results showed no age-related changes in any of our regions of interest (Figure 2; Supplementary Table 3). However, results showed a significant age-by-hemisphere interaction in the AIns (*β* = -0.39, *p* = 5.88×10^−5^), which was driven by age-related increase in Glu/GABA ratio in the right AIns (*β* = -0.28, *p* = 0.00043) that was not significant in the left hemisphere (*β* = 0.11, *p* = 0.11).

**Figure 2:**
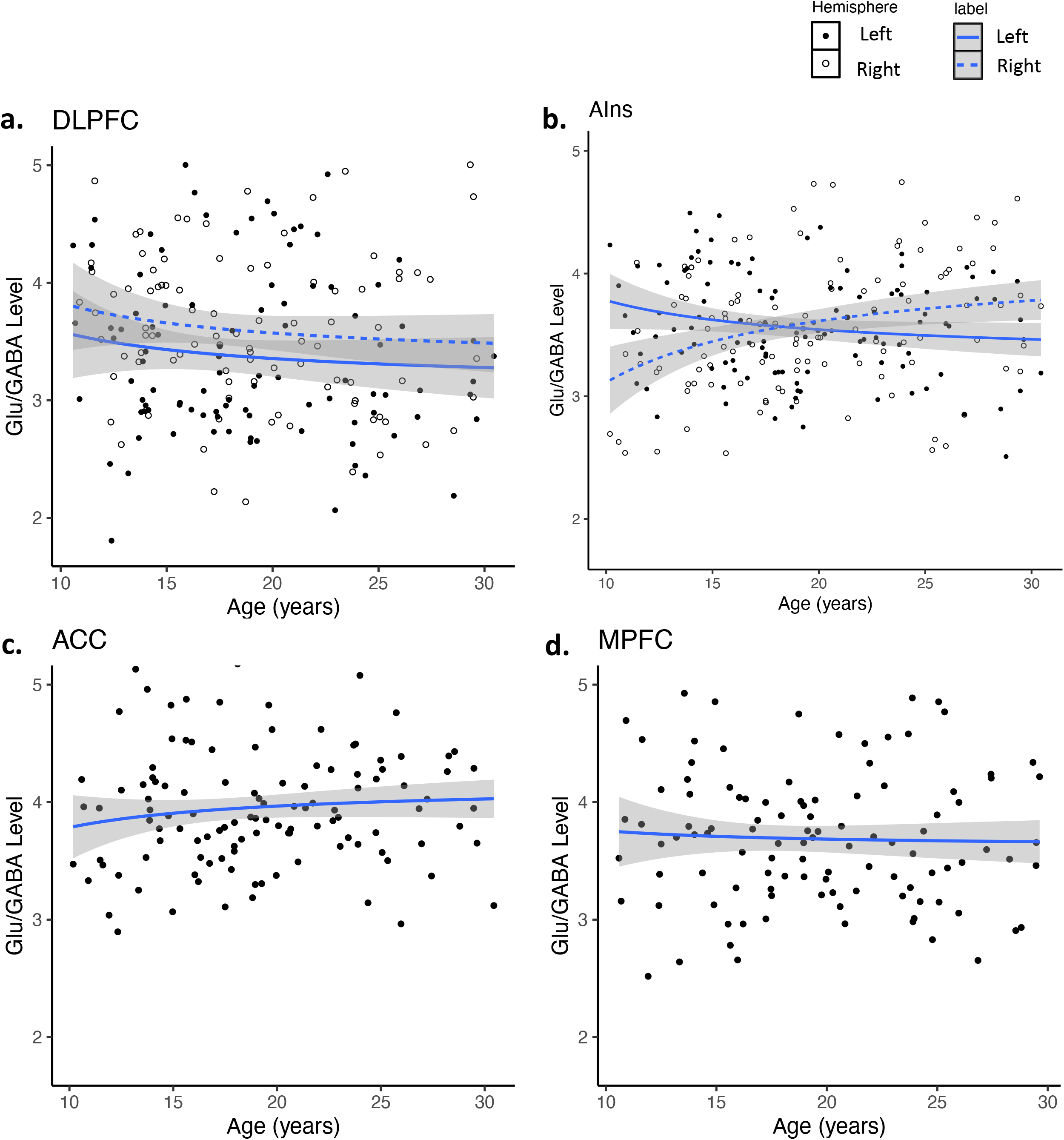
Glutamate/GABA age effects across PFC regions of interest. Statistical values reflect linear mixed effects model fits for regions with both a left and right hemisphere and linear model fits for midline regions. All are modeled with an inverse function. Solid line and dots reflect left hemisphere and dashed line and open dots reflect right hemisphere where applicable. Legend refers to figures (A) and (B). (A) left and right dorsolateral prefrontal cortex (B) left and right anterior insula (C) midline anterior cingulate cortex (d) midline medial prefrontal cortex.

### Decreased inter-subject variability through adolescence

Age-related changes in inter-subject variability in metabolite levels were assessed using models of age-related change (described above) with the best fitting functional form of age (i.e., inverse age), and computing the absolute value of the residual of each metabolite relative. This approach captures the absolute deviation from the age-adjusted mean, with higher values indicating greater deviation, and lower values indicating metabolite levels very near the expected level for the participant’s age. Absolute deviation values were then assessed as a function of age to measure inter-subject variability. Due to the main effect of hemisphere as well as interaction effect, hemispheres were considered separately for all further analyses. Age-related decreases in variability in Glu/Cr were identified in ACC (*β* = 0.26, *p* = 0.018), right AIns (*β* = 0.24, *p* = 0.039), MPFC (*β* = 0.26, *p* = 0.021), and left DLPFC (*β* = 0.32, *p* = 0.0026) (Figure 3) but not in left AIns or right DLPFC (Supplementary Table 4). Inter-subject variability in GABA/Cr did not change with age in any ROI (Figure 4; Supplementary Table 5).

**Figure 3:**
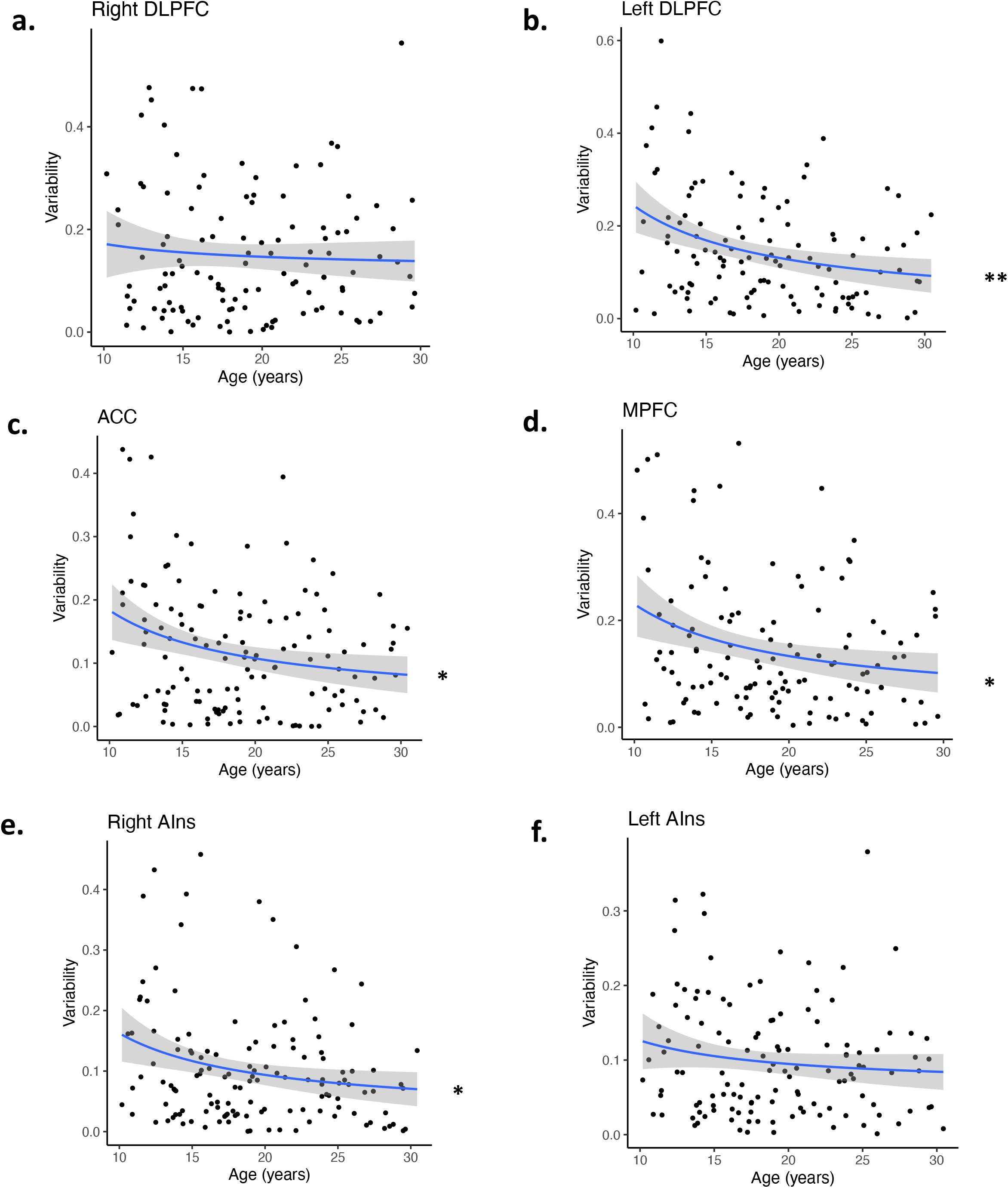
Age effects of inter-individual variability of glutamate/Cr. Statistical values reflect linear model fits for midline regions. All are modeled with an inverse function. (A) right dorsolateral prefrontal cortex (B) left dorsolateral prefrontal cortex (C) midline anterior cingulate cortex (D) midline medial prefrontal cortex (E) right anterior insula (F) left anterior insula. * *p* < .05, ** *p* < .01, *** *p* < .001. Asterisks reflect Bonferroni-corrected p-values.

**Figure 4:**
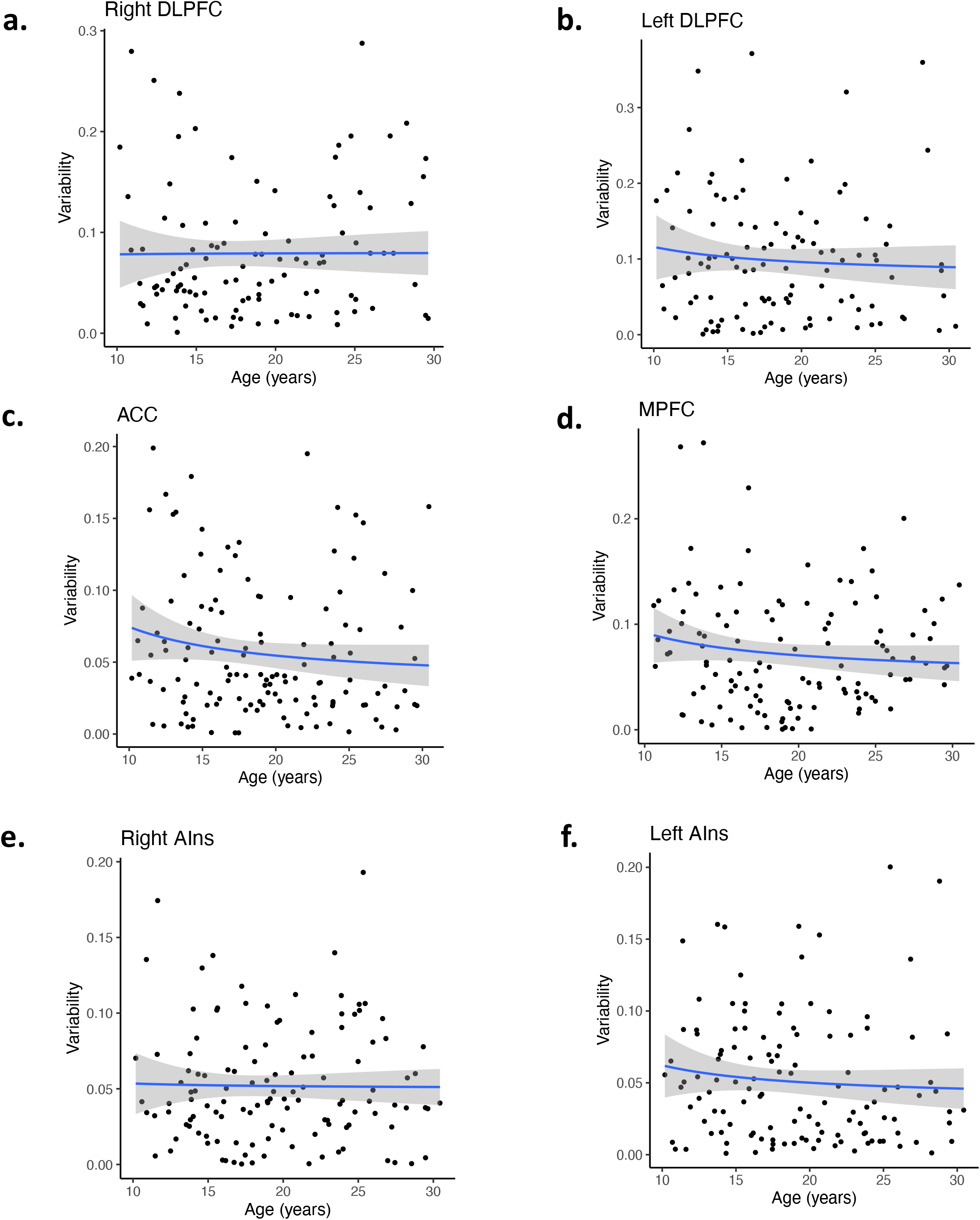
Age effects of inter-individual variability of GABA/Cr. Statistical values reflect linear model fits for midline regions. All are modeled with an inverse function. (A) right dorsolateral prefrontal cortex (B) left dorsolateral prefrontal cortex (C) midline anterior cingulate cortex (D) midline medial prefrontal cortex (E) right anterior insula (F) left anterior insula. \**p* < .05, ** *p* < .01, *** *p* < .001. Asterisks reflect Bonferroni-corrected p-values.

### Emergence of E/I balance with age

To determine the extent to which age-related changes in GABA/Cr and Glu/Cr are inter-related, we measured GABA/Cr-Glu/Cr correlation as a function of age based on a previously reported approach (Steel et al., 2020). While the Glu/GABA ratio provides a measure of the relative over- or under-abundance of each metabolite, the correlation provides a measure of how glutamate and GABA levels tend to covary by region (Rideaux, 2021; Steel et al., 2020). That is, if Glu and GABA are balanced, then the relative level of GABA should be associated with the relative level of Glu (i.e., both high, or both low), whereas relative mismatches in GABA and Glu levels (e.g., high GABA but low Glu, or vice versa) would be indicative of Glu-GABA imbalance. In order to examine whether Glu/Cr and GABA/Cr were correlated within regions in our sample, and to what extent these correlations varied with age, we used linear regression to examine the relationship between GABA/Cr and Glu/Cr and their interaction with age. Results showed that Glu/Cr and GABA/Cr were significantly associated in all our ROIs (Supplementary Table 6) and this balance increased with age in the right AIns (*β* = -0.20, *p* = 0.042), ACC (*β* = -0.22, *p* = 0.0102), MPFC (*β* = -0.19, *p* = 0.037), and a trend-level interaction in the right DLPFC (*β* = -0.21, *p* = 0.06) (Supplementary Table 7). There were no age-related effects in L DLPFC (*β* = -0.08, *p* = 1.00) or L AIns (*β* = -0.11, *p* = 1.00).To visualize these interactions non-parametrically, we computed group-wise correlations between GABA and Glu (Figure 5a), and used a sliding window approach to assess the effect size of Glu-GABA associations within each age bin (Figure 5b). Both approaches illustrate that significant interactions reflect age-related increases Glu-GABA balance.

**Figure 5:**
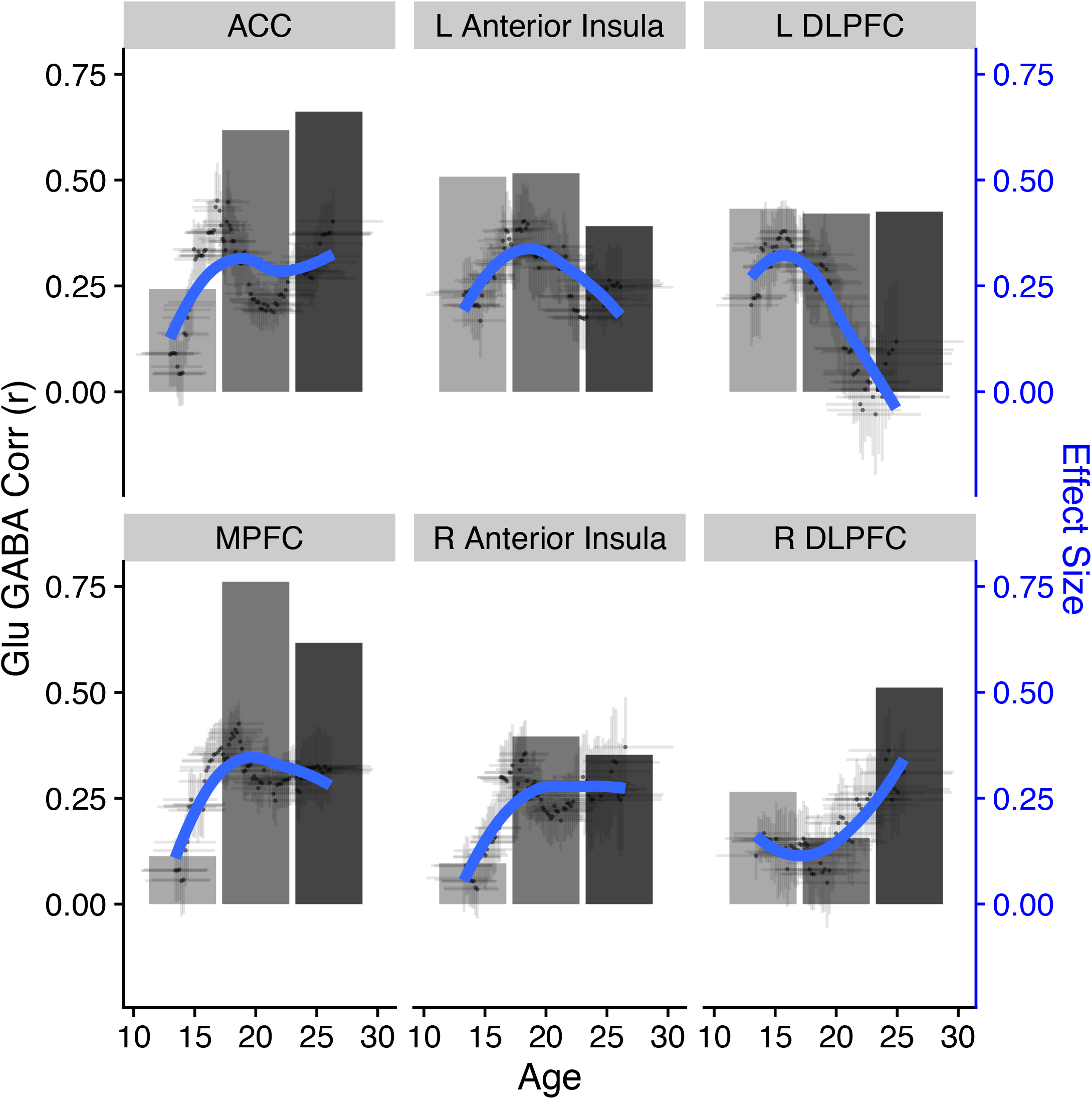
Association between glutamate/Cr and GABA/Cr by age. Panels show association for each ROI (top: anterior cingulate cortex, left anterior insula, left dorsolateral prefrontal cortex; bottom: medial prefrontal cortex, right anterior insula, right dorsolateral prefrontal cortex). Grayscale bars (left y-axis) reflect correlation (r value) between GABA/Cr and Glu/Cr within each age group. Blue lines (right y-axis) reflect effect sizes of the GABA/Cr-Glu/Cr regression coefficients computed within a 30-subject sliding window (individual dots represent the association within each sliding window, plotted at the mean age within the window, and with x-error bars indicating the age range span, and y-error bars indicating the effect size SEM; blue lines are a loess curve fit across age windows).

## Discussion

This study describes a novel high-field (7T) MR spectroscopic imaging approach using a large dataset to characterize age-related changes in glutamate/Cr and GABA/Cr in PFC to identify markers of critical period plasticity in adolescence. In accordance with our hypotheses, we found evidence for global age-related decreases in Glu/Cr across PFC. Contrary to our hypotheses, we did not find evidence for increases in GABA/Cr through adolescence, which appeared stable in DLPFC and MPFC, while showing decreases in AIns and ACC. Importantly, while Glu/GABA ratios did not show systematic age-related change, we found age-related increases in the correlation of these metabolites suggesting E/I becoming balanced into adulthood. Overall, these results, when coupled with evidence from the extant animal and postmortem literature, provide compelling new evidence that through adolescence, there is dynamic change in excitatory and inhibitory balance informing models of an adolescent critical period of plasticity (Larsen et al., 2021).

MRSI evidence of decreases in glutamate through adolescence may be driven by known related brain maturational changes including decreases in the density of pyramidal cell dendritic spines, particularly in layer 3 prefrontal cortex (Bourgeois et al., 1994; Huttenlocher, 1999; Petanjek et al., 2011; Rakic et al., 1986), reductions in AMPA GluR1 subunits (Datta et al., 2015; Hoftman et al., 2017), NMDAR NR3A subunits (Henson et al., 2008), and genes involved in glutamate signaling (Harris et al., 2009). Indeed, a recent multimodal study found significant associations between MRSI Glu/Cr and PET-based measures of synaptic density using the radioligand [11C]UCB-J (Onwordi et al., 2021), supporting that changes in Glu may be associated with underlying developmental mechanisms such as synaptic pruning.

Additionally, many of the regions in which we observed significant age-related decreases in metabolite levels showed age-related decreases in inter-subject variability of Glu/Cr. Inter-subject variability may be driven by greater intra-subject variability, as is evident in behavior (Klein et al., 2011; McIntosh et al., 2008; Montez et al., 2017; Tamnes et al., 2012) and brain function during cognitive tasks (Montez et al., 2017). Developmental decreases in variability suggest that glutamate levels may fluctuate through adolescence, reflecting an adaptive period of plasticity that may be necessary for stabilization in adulthood. This is supported by our findings showing greater correlation between Glu/Cr and GABA/Cr in adulthood, reflecting convergence and equifinality with maturation. Future longitudinal follow-up studies delineating individual trajectories can more directly inform this hypothesis.

There is strong evidence for increases in GABAergic transmission from animal models and post-mortem human studies, specifically in aspects of inhibition related to PV neurons. However, we did not find any regions exhibiting age-related increases in GABA. While this was in contrast to our hypotheses, it is notable there are other aspects of PFC GABA that show divergent age-related trajectories through adolescence, including stability of GABAergic synapse density (Bourgeois et al.,1994), decreases in PV positive cartridges (Cruz et al., 2003) and GABA alpha 2 subunit expression (Hashimoto et al., 2009), and increases in PV positive varicosities (Erickson and Lewis 2002). Taken together, these results could indicate that while MRS is not sensitive enough to pick up on specific changes in PV-positive neurons, it instead reflects a composite view of many aspects of GABA signaling, providing a more generalized measure of the overall tone of GABAergic inhibition.

Instead, we found that GABA/Cr exhibited a more regionally heterogeneous pattern of age effects with no change in DLPFC and MPFC, and decreases in AIns and ACC. These divergent developmental trajectories may reflect known differences in the functional recruitment of PFC through adolescence. For example, the activation of the ACC has been found to increase through adolescence into adulthood, mediating development of inhibitory control, while the AIns exhibits a protracted structural and functional developmental trajectory and is associated with attentional and interoceptive processes, aspects of cognition that show late development through adolescence (Uddin et al., 2011). This is in contrast to the DLPFC, which has been found to be engaged at adult-level levels by adolescence during cognitive control tasks (Ordaz et al., 2013; Simmonds et al., 2017). Consistent with this view, we found that the AIns and ACC showed protracted age-related decreases in GABA/Cr through our adolescent sample, while DLPFC and

MPFC were already at adult levels. GABA decreases in the AIns and ACC may reflect known decreases in calretinin neurons and PV-containing chandelier cells (Caballero et al., 2014; Fish et al., 2013). Notably, evidence suggests that GABA measured with MRS may reflect tonic neurotransmitter function instead of phasic processes (Dyke et al., 2017). Consistent with our results, the size of tonic inhibitory currents has been shown to decline through adolescence in mouse cingulate cortex (Piekarski et al., 2017). In contrast, the lack of age effects for GABA levels in the DLPFC and MPFC are in line with a recent meta-analysis of MRS data showing GABA stability through adolescence (Porges et al., 2021). This could reflect either an earlier developmental plateau, consistent with functional data, or opposing GABAergic processes, such as increases in PV and decreases in calretinin, that may give the appearance of stability in our MRSI data.

Given the substantial differences between 3T single-voxel MRS and our 7T multi-voxel MRSI data, it is likely not surprising that our findings diverge from several earlier studies. 3T MRI studies have typically not found age related changes in glutamate (Blüml et al., 2013; Ghisleni et al., 2015; Gleich et al., 2015; Silveri et al., 2013), except for (Shimizu et al., 2017). One key difference from these previous studies is that 7T MRSI allows us to separate the overlapping signals of glutamate and glutamine, and thus the decreases we observe may be more specifically associated with Glu, rather than the combined signal typically reported at 3T. Further, some 3T studies also show increases in GABA in adults compared to adolescents (Ghisleni et al., 2015; Silveri et al., 2013). However, lower-field strength GABA data is confounded by overlapping macromolecule signal that can have significant impacts on results (Bell et al., 2021). Importantly, our 7T voxels are 9 mm by 9 mm by 10 mm as compared to 3T MRS voxels, which are frequently larger. This allows us to more precisely define specific ROIs, as well as prioritize a higher fraction of gray matter in the voxel, all of which could have an impact on metabolite measurement (Choi et al., 2006). However, even at 7T, MRSI provides a measurement of total metabolite levels within a voxel, and although we can provide greater specificity in metabolite measurement, we cannot differentiate the unique contributions of that metabolite in neurotransmission from involvement in other biological or metabolic processes.

Glutamate/GABA balance has been typically assessed by calculating their ratio, assuming that they are positively related. We did not find age related changes in the ratio of glutamate and GABA. The lack of widespread age-related changes in the Glu/GABA ratio indicates that aggregate metabolite levels do not show systematic age-related shifts within this age range. Substantial individual variability in adolescence in these metabolites may contribute to differences in their correlation across ages, rendering the ratio less likely to identify changes beyond those indicated by the individual metabolites alone (Steel et al., 2020). Indeed, our results show that Glu/Cr and GABA/Cr become more correlated with age; that is, those participants with higher or lower glutamate relative to the group also tended to have correspondingly higher or lower GABA, which may reflect greater E/I balance, as has been found in parietal cortex (Steel et al., 2020).

Together, these findings provide compelling new evidence of increases in E/I balance through adolescence in PFC that may be consistent with critical period plasticity. Notably, cognitive development proceeds in a similar timeline to the developmental changes reported here, with improvements in executive function through adolescence into adulthood (Ordaz et al., 2013; Simmonds et al., 2017). Greater glutamate/GABA balance may support cognitive functions by enhancing signal-to-noise ratio and synchrony of neural activity (Larsen & Luna, 2018). Understanding the brain mechanisms underlying the transition from adolescence to adulthood informs our understanding of the processes driving normative development. This is critical for understanding disorders which may emerge as a result of impaired E/I balance developmental trajectories as schizophrenia, which often first emerge in adolescence and has been associated with alterations in the balance of GABA and glutamate (Glantz LA & Lewis DA, 2000; Gonzalez-Burgos & Lewis, 2008; Hoftman & Lewis, 2011). Gaining a more complete understanding of impaired developmental trajectories in GABA and glutamate could better inform much-needed pharmaceutical therapies that act on these neurotransmitters. Future longitudinal studies are needed to determine individual developmental trajectories, and to examine associations with behavioral indices of cognitive development as well as indices of improvements in signal-to-noise ratio.

## Methods

### Participants

MRSI data were collected on 144 10–30 years old participants (73 females, see Supplementary Figure 1) recruited from the community. Participants were excluded if they had a history of head injury with loss of consciousness, a non-correctable vision problem, a history of substance abuse, a learning disability, a history of major psychiatric or neurologic conditions in themselves or a first-degree relative. Participants were also excluded if they reported any MRI contraindications, such as non-removable metal in their body. For participants under the age of 18, parental consent and participant assent were obtained prior to data collection. For participants over the age of 18, participant consent was obtained. Participants received payment for their participation. All experimental procedures were approved by the University of Pittsburgh Institutional Review Board and complied with the Code of Ethics of the World Medical Association (Declaration of Helsinki, 1964).

### MRSI Data Acquisition

This study was performed at the University of Pittsburgh Medical Center Magnetic Resonance Research Center in a Siemens 7T scanner. Structural images were acquired using an MP2RAGE sequence (1 mm isotropic resolution, TR/TE/flip angle 1/ flip angle 2: 6000 ms/2.47 ms/40/50) for parcellation and alignment. MRSI of GABA and glutamate were acquired using a J-refocused spectroscopic imaging sequence (Pan et al., 2010) (TE/TR = 2×17/1500ms) in order to minimize the impact of momentary interactions between separate molecules on chemical shifts. Radiofrequency (RF) based outer volume suppression was used to minimize interference in the signal from extracerebral tissue (Hetherington et al., 2010). An 8×2 1H transceiver array using 8 independent RF channels was used to acquire data. High order shimming was used to optimize homogeneity of the B0 magnetic field.

One oblique axial slice (10mm thick, 24×24 encoded across a FOV of 216×216 mm, 0.9×0.9×1.0 cm effective resolution) was acquired and positioned to ensure that the DLPFC (roughly defined as Brodmann Area 9) was present (Figure 6a). MRSI slice positioning was accomplished by aligning the T1 anatomical images of each participant to MNI space, and then aligning the MRSI acquisition to the anatomical image. An in-house program - Quantitative Partial Acquisition Slice-Alignment (Q-Pasa) - was used to ensure equivalent slice acquisitions across individual variability in brain morphology and slice positioning. Q-Pasa maps an oblique slice atlas (standard MNI space) into a participant’s native space during the scan using BA9 and the thalamus as reference points in order to guide placement of the acquired slice.

**Figure 6:**
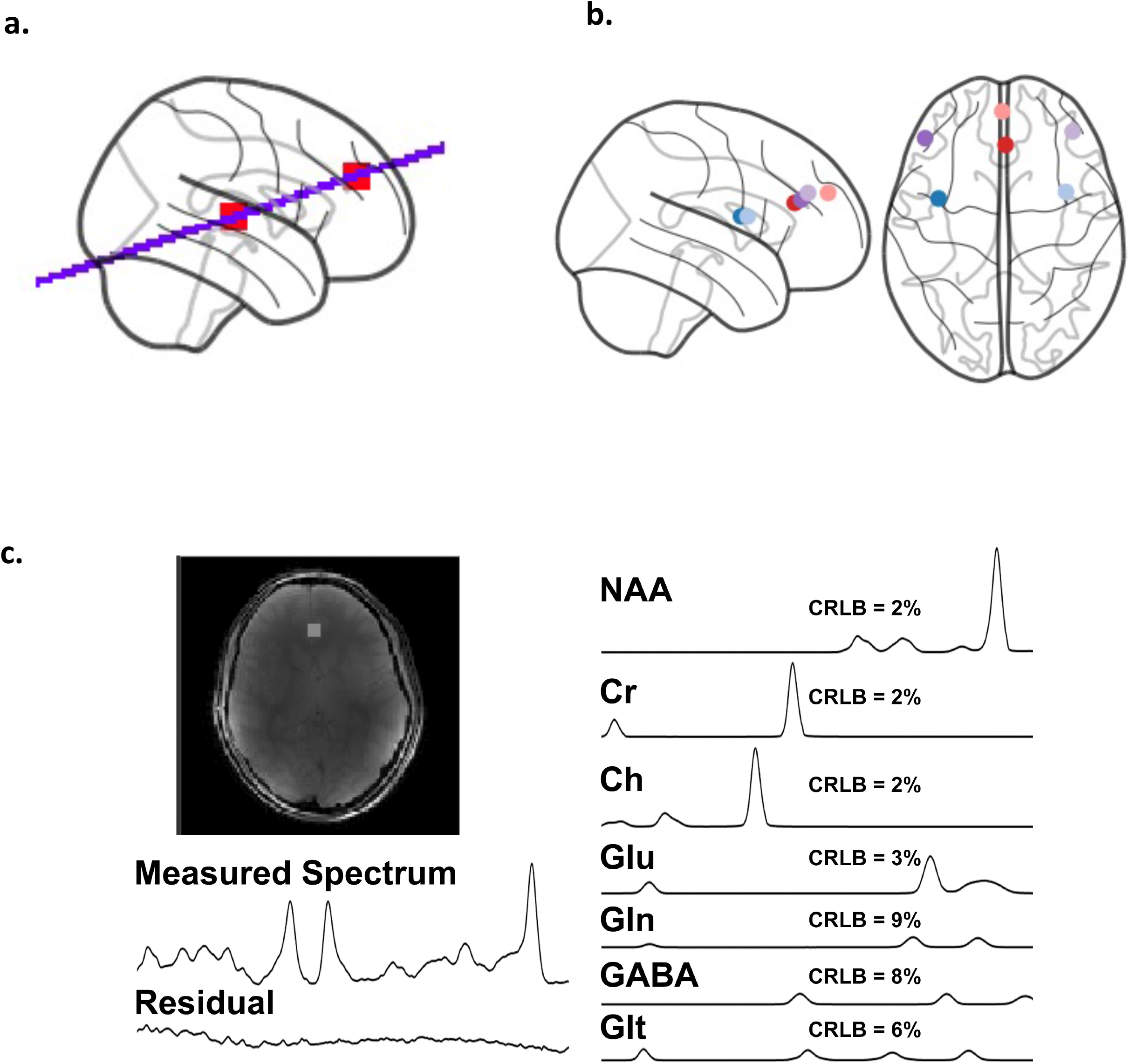
(a) ideal oblique MRSI slice position through DLPFC and thalamus (shown in red boxes). (b) Typical positioning of all assessed ROIs (MPFC = salmon; ACC = red; right and left DLPFC = purple; right and left Anterior Insula = blue) (generated with code adapted from Westwater et al., 2022) (c) Representative spectral data from a typical participant’s ACC ROI. Measured spectrum shows a typical acquired signal, and residual indicates the difference between the raw data and the combined signals of all fit metabolite spectra (shown right with CRLB estimates).

### Metabolite Levels

Quantification of neurotransmitters was achieved by fitting MRSI data using LCModel (Provencher, 2001). The LCModel output provides estimates of metabolite levels for all metabolites present, including metabolites of interest such as Glutamate and GABA, as well as major metabolites like glycerophosphocholine + phosphocholine + choline (GPC+Cho), N-Acetylaspartate + N-Acetylaspartylglutamic Acid (NAA+NAAG), and creatine (Cr). LCModel outputs contain Cramer-Rao Lower Bounds (CRLB), which provide an expression of the uncertainty of the estimate for each metabolite. The CRLB provides an estimate of the lowest achievable variance for an estimator and is used as an indicator of how well the measurement of a given metabolite fits the ideal spectrum for that metabolite, providing an assessment of data quality. Additionally, outputs were used to visually inspect spectra and model fit, which we used as a first-pass exclusion criteria for data, along with the CRLB.

Brain neurotransmitter levels are given as metabolite ratios of the neurotransmitter of interest to creatine (Cr) (Glu/Cr or GABA/Cr). This allows for a shorter acquisition time (as compared to using a water reference) while also providing for a means to control for inter-subject variability due to the factors such as an amount of cerebrospinal fluid in the voxel. Cr is used as the denominator in metabolite ratios because it is a metabolite with a strong signal and reliable chemical shift, in addition to being relatively stable under healthy conditions (Rackayova et al., 2017) and with age (Saunders et al., 1999).

### ROI-Based Analysis

Regions of interest (ROIs) were defined in MNI-space using a custom ROI atlas, and included right and left dorsolateral prefrontal cortex (DLPFC), right and left anterior insula (AI), anterior cingulate cortex (ACC), and medial PFC (MPFC) (Figure 6b). An automated nonlinear registration using FSL’s FNIRT was performed to map those ROIs into each subject’s native space. ROI centers were visually inspected and manually adjusted for each subject with in-house developed software in order to maximize spectral quality and gray matter content of the voxel. To ensure manual adjustment did not move ROI centers outside of the region of interest, the adjusted ROIs were registered back to standard space where their position was confirmed with the Eickhoff-Zielles maximum probability map of the Talairach-Tournoux atlas using AFNI’s whereami (Cox, 1996). The ROI coordinates were then used in the anatomically guided voxel-shifting reconstruction of MRSI data (Hetherington et al., 2007) to gather spectroscopic data over the ROI.

### Data Quality Criteria

LCModel fits were first visually inspected, and data were excluded for bad model fits due to obvious artifacts, lipid contamination or baseline distortion. Data were then excluded if any of the three major metabolite peaks, GPC/Cho, NAA/NAAG, and Cr had a CRLB greater than 10% or if the macromolecule/Cr content in their spectra was greater than 3, in order to remove distorted data that would lead to improper estimates of harder to detect metabolites. Consistent with generally accepted practice, GABA/Cr and Glu/Cr data were excluded if the CRLB of GABA or glutamate, respectively, was greater than 20. Data were excluded at the ROI level, so participants that had poor data in one region were not necessarily excluded from all analyses if they had high quality data in another region. For group level analyses assessing the relationship between metabolite levels and age, a threshold of 2 standard deviations from the mean was used to exclude outliers. For the analysis looking at the association between glutamate and GABA, only participants that had both good GABA and good glutamate data (as defined by the CRLB cutoff) were included. A representative spectrum is shown in Figure 6c.

None of the regions tested showed worse data quality (higher CRLB) among younger subjects (Supplementary Table 8). One region (MPFC) showed higher CRLB among older subjects, but this is unlikely to account for the effect reported as those with the highest correlations had worse data quality.

### Statistical Analysis

To examine age-related change in metabolite levels, linear mixed-effects regression analyses were implemented through the R packages lme4 and lmerTest (Bates et al., 2014; Kuznetsova et al., 2017). For regions that had both a left and a right hemisphere (AIns and DLPFC), age-related changes in neurotransmitter measures were examined using linear mixed-effects models with participant as a random effect and ROI hemisphere, sex, and fraction of gray matter in the voxel as covariates. For midline regions (the ACC and MPFC), hemispheres were not considered and linear regression was used, with sex and fraction of gray matter as covariates. All models were tested with and without these covariates.

Three forms of age effects were tested: linear age, inverse age, and quadratic age; these are the functional forms that have previously characterized age-related change during this developmental period (Luna et al., 2004). The best form of age was chosen using Akaike’s Information Criterion (AIC), wherein the model with the lowest AIC was chosen as the best fit model (Akaike, 1974). For the majority of models, inverse age was the best fit (Supplementary Tables 9, 10, 11). For models in which no form of age was significantly different from the others, inverse age was chosen for consistency. To correct for multiple comparisons, we report adjusted p-values based on Bonferroni-correction, using the R package ‘stats’ (R Core Team 2013), and a correction based on the number of ROIs being tested. Variables were z-scored to obtain standardized coefficients.

To examine age-related changes in inter-subject variability, we used linear regression to test for age effects in the absolute residuals (i.e., deviation from the age-adjusted mean) of metabolite levels. Given the age-by-hemisphere interaction in the Glu/GABA ratio of the AI, we considered all regions separately for this analysis. Inverse age was used as that was previously characterized as the best model for both GABA and glutamate individually.

In addition to metabolite levels, we were interested in examining correlation among metabolites, based on recent studies suggesting a balance of GABA and glutamate among young adult subjects (Steel et al., 2020). To examine age-related changes in metabolite correlation, we used linear regression to look at the association between GABA and Glutamate and their interaction with age across the sample within each region individually while controlling for the same covariates as before. The interaction was visualized in two ways. First, correlation coefficients (r values) were computed for each age group (10-16; 17-22; 23-30) of roughly equal sample size. Covariates were first residualized out prior to computing the correlation. Second, slope and standard error are modeled across a sliding window (each with 30 participants) while controlling for covariates.

