## Supplementary Figures & Tables for "Changes in prefrontal GABA and glutamate through adolescence supports excitation/inhibition balance"

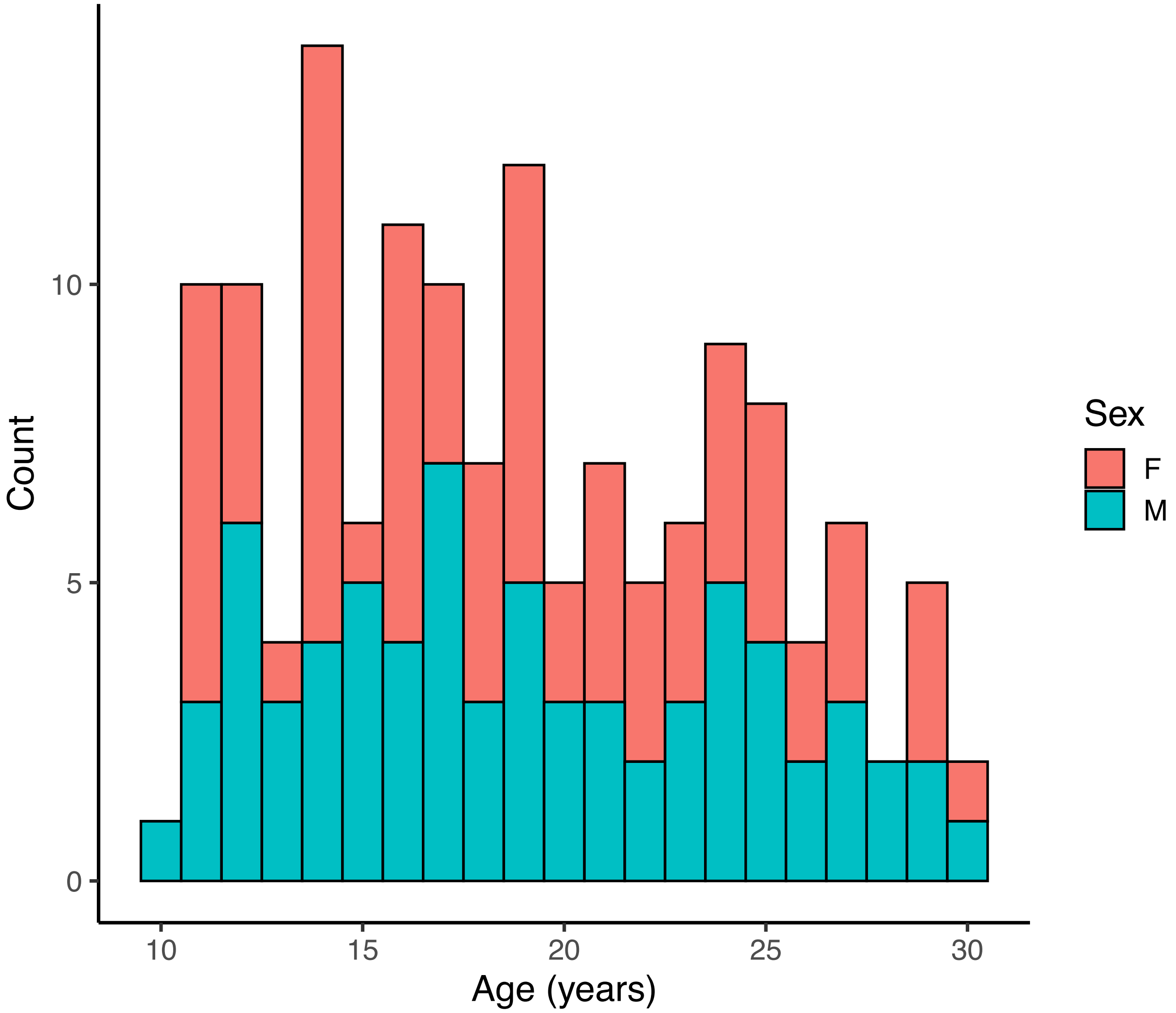


Supplementary Figure 1: participant age histogram

Supplementary Tables

Supplementary Table 1: Glu/Cr model output for all regions

| **ROI** | **Variable** | **β** | **t** | **n participants** | **P (uncorrected)** | **P (Bonferroni corrected)** |
|---|---|---|---|---|---|---|
| L & R DLPFC | **Age$^{-1}$** | 0.15 | 2.81 | 131 | 0.006 | 0.024 |
| | Sex | 0.04 | 0.41 | 131 | 0.68 | 1.00 |
| | **Hemisphere** | -0.40 | -5.01 | 131 | $1.96 \times 10^{-6}$ | $7.84 \times 10^{-6}$ |
| | Frac GM | 0.03 | 0.64 | 131 | 0.52 | 1.00 |
| ACC | **Age$^{-1}$** | 0.21 | 3.248 | 132 | 0.0015 | 0.006 |
| | Sex | 0.02 | 0.15 | 132 | 0.88 | 1.00 |
| | Frac GM | -0.014 | -0.21 | 132 | 0.84 | 1.00 |
| MPFC | Age$^{-1}$ | 0.11 | 1.70 | 124 | 0.093 | 0.37 |
| | Sex | 0.11 | 0.84 | 124 | 0.40 | 1.00 |
| | Frac GM | 0.078 | 1.24 | 124 | 0.22 | 0.88 |
| L & R AIns | **Age$^{-1}$** | 0.29 | 6.33 | 136 | $3.29\text{x}10^{-9}$ | $1.3129\text{x}10^{-8}$ |
| | Sex | 0.09 | 1.09 | 136 | 0.28 | 1.00 |
| | **Hemisphere** | -0.76 | -14.89 | 136 | $5.51\text{x}10^{-29}$ | $2.20\text{x}10^{-28}$ |
| | Frac GM | 0.06 | 1.68 | 136 | 0.09 | 0.36 |

Supplementary Table 2: GABA/Cr model output for all regions

| **ROI** | **Variable** | **β** | **t** | **n participants** | **P (uncorrected)** | **P (Bonferroni corrected)** |
|---|---|---|---|---|---|---|
| L & R DLPFC | Age$^{-1}$ | -0.03 | -0.64 | 128 | 0.52 | 1.00 |
| | Sex | -0.16 | -1.60 | 128 | 0.11 | 0.44 |
| | **Hemisphere** | -0.26 | -3.31 | 128 | 0.001 | 0.004 |
| | **Frac GM** | 0.14 | 3.17 | 128 | 0.002 | 0.008 |
| ACC | **Age$^{-1}$** | 0.27 | -2.01 | 130 | $6.38\text{x}10^{-5}$ | $2.55 \text{x}10^{-4}$ |
| | Sex | 0.12 | 0.97 | 130 | 0.34 | 1.00 |
| | Frac GM | -0.03 | -0.47 | 130 | 0.64 | 1.00 |
| MPFC | Age$^{-1}$ | 0.10 | 1.53 | 120 | 0.13 | 0.52 |
| | Sex | 0.002 | 0.02 | 120 | 0.98 | 1.0 |
| | Frac GM | 0.08 | 1.29 | 120 | 0.20 | 0.80 |
| L & R AIns | **Age$^{-1}$** | 0.22 | 4.77 | 128 | $4.71\text{x}10^{-6}$ | $1.88\text{x}10^{-5}$ |
| | Sex | -0.08 | -0.95 | 135 | 0.34 | 1.00 |
| | **Hemisphere** | -0.49 | -6.05 | 135 | $1.64 \text{x}10^{-8}$ | $6.56\text{x}10^{-8}$ |
| | Frac GM | -0.11 | -2.46 | 135 | 0.015 | 0.06 |
| | Age$^{-1}$ x Hemisphere | 0.16 | 1.99 | 135 | 0.048 | 0.192 |

Supplementary Table 3: Glutamate/GABA ratio model output for all regions

| ROI | Variable | β | t | n participants | P (uncorrected) | P (Bonferroni corrected) |
|---|---|---|---|---|---|---|
| R & L DLPFC | $Age^{-1}$ | 0.09 | 1.45 | 117 | 0.15 | 0.60 |
| | Sex | 0.07 | 0.65 | 117 | 0.52 | 1.00 |
| | Hemisphere | 0.20 | 2.17 | 117 | 0.04 | 0.16 |
| | Frac GM | -0.09 | -1.73 | 117 | 0.08 | 0.32 |
| ACC | $Age^{-1}$ | -0.09 | -1.31 | 117 | 0.19 | 0.76 |
| | Sex | -0.12 | -0.90 | 117 | 0.37 | 1.00 |
| | Frac GM | -0.018 | -0.28 | 117 | 0.78 | 1.00 |
| | $Age^{-1}$ x Sex | -0.27 | -2.04 | 117 | 0.043 | 0.17 |
| MPFC | $Age^{-1}$ | 0.04 | 0.44 | 107 | 0.66 | 1.00 |
| | Sex | 0.013 | 0.075 | 107 | 0.94 | 1.00 |
| | Frac GM | -0.04 | -0.42 | 107 | 0.68 | 1.00 |
| L & R AIns | $Age^{-1}$ | -0.09 | -1.75 | 131 | 0.13 | 0.52 |
| | Sex | 0.12 | 1.10 | 131 | 0.10 | 0.40 |
| | Hemisphere | 0.02 | 0.22 | 131 | 0.83 | 1.00 |
| | Frac GM | 0.14 | 2.60 | 131 | 0.011 | 0.044 |
| | **$Age^{-1}$ x Hemisphere** | -0.39 | -4.54 | 131 | $1.47x10^{-5}$ | $5.88x10^{-5}$ |

Supplementary Table 4: Inter-individual variability of Glu/Cr

| ROI | Variable | β | t | n participants | P (uncorrected) | P (Bonferroni corrected) |
|---|---|---|---|---|---|---|
| L DLPFC | $Age^{-1}$ | 0.06 | 0.68 | 116 | 0.50 | 1.00 |
| | Frac GM | -0.02 | -0.23 | 116 | 0.82 | 1.00 |
| | Sex | -0.18 | -0.97 | 116 | 0.34 | 1.00 |
| R DLPFC | **$Age^{-1}$** | 0.29 | 3.28 | 120 | 0.0014 | 0.0084 |
| | Frac GM | 0.23 | 2.59 | 120 | 0.011 | 0.066 |
| | Sex | -0.051 | -0.29 | 120 | 0.77 | 1.00 |
| ACC | **$Age^{-1}$** | 0.27 | 3.10 | 133 | 0.0024 | 0.0144 |
| | Frac GM | -0.01 | -0.12 | 133 | 0.91 | 1.00 |
| | Sex | 0.12 | 0.72 | 133 | 0.48 | 1.00 |
| MPFC | **$Age^{-1}$** | 0.27 | 2.99 | 125 | 0.0034 | 0.0204 |
| | Frac GM | -0.04 | -0.50 | 125 | 0.62 | 1.00 |
| | Sex | -0.26 | -1.51 | 125 | 0.13 | 0.78 |
| L AIns | $Age^{-1}$ | 0.10 | 1.11 | 127 | 0.27 | 1.00 |
| | Frac GM | 0.12 | 1.42 | 127 | 0.16 | 0.96 |
| | Sex | -0.18 | -1.03 | 127 | 0.31 | 1.00 |
| R AIns | $Age^{-1}$ | 0.21 | 2.4 | 134 | 0.017 | 0.10 |

| | | | | | | |
|---|---|---|---|---|---|---|
| | Frac GM | 0.13 | 1.47 | 134 | 0.14 | 0.84 |
| | Sex | -0.19 | 1.12 | 134 | 0.27 | 1.00 |

Supplementary Table 5: Inter-individual variability of GABA/Cr

| **ROI** | **Variable** | **β** | **t** | **n** **participants** | **P (uncorrected)** | **P (Bonferroni corrected)** |
|---|---|---|---|---|---|---|
| L DLPFC | $Age^{-1}$ | 0.11 | 1.09 | 110 | 0.28 | 1.00 |
| | Frac GM | -0.07 | -0.73 | 110 | 0.47 | 1.00 |
| | Sex | 0.21 | 1.10 | 110 | 0.28 | 1.00 |
| R DLPFC | $Age^{-1}$ | -0.017 | -0.18 | 107 | 0.86 | 1.00 |
| | Frac GM | 0.19 | 1.94 | 107 | 0.06 | 0.36 |
| | Sex | 0.16 | 0.79 | 107 | 0.43 | 1.00 |
| ACC | $Age^{-1}$ | 0.14 | 1.61 | 131 | 0.11 | 0.66 |
| | Frac GM | -0.22 | -2.69 | 131 | 0.0082 | 0.050 |
| | Sex | -0.36 | -2.13 | 131 | 0.035 | 0.21 |
| MPFC | $Age^{-1}$ | 0.14 | 1.39 | 121 | 0.17 | 1.00 |
| | Frac GM | -0.062 | -0.68 | 121 | 0.50 | 1.00 |
| | Sex | -0.28 | -1.56 | 121 | 0.12 | 0.72 |
| L AIns | $Age^{-1}$ | 0.11 | 1.14 | 122 | 0.26 | 1.00 |
| | Frac GM | -0.13 | -1.37 | 122 | 0.17 | 1.00 |
| | Sex | -0.33 | -1.82 | 122 | 0.071 | 0.43 |
| R AIns | $Age^{-1}$ | -0.003 | -0.033 | 125 | 0.97 | 1.00 |
| | Frac GM | 0.011 | 0.12 | 125 | 0.91 | 1.00 |
| | Sex | -0.19 | -1.044 | 125 | 0.30 | 1.00 |

Supplementary Table 6: Model output of main effects of the association between GABA/Cr and Glu/Cr

| **ROI** | **Variable** | **β** | **t** | **n** **participants** | **P (uncorrected)** | **P (Bonferroni corrected)** |
|---|---|---|---|---|---|---|
| L DLPFC | **Glu/Cr** | 0.40 | 4.33 | 102 | $3.49x10^{-5}$ | $2.1x10^{-4}$ |
| R DLPFC | **Glu/Cr** | 0.32 | 3.38 | 100 | 0.001 | 0.006 |
| ACC | **Glu/Cr** | 0.51 | 6.44 | 123 | $2.58x10^{-9}$ | $1.55x10^{-8}$ |
| MPFC | **Glu/Cr** | 0.48 | 5.78 | 113 | $7.04x10^{-8}$ | $4.22x10^{-7}$ |
| L AIns | **Glu/Cr** | 0.50 | 6.12 | 116 | $1.39x10^{-8}$ | $8.34x10^{-8}$ |
| R AIns | **Glu/Cr** | 0.38 | 4.52 | 124 | $1.46x10^{-5}$ | $8.76x10^{-5}$ |

Supplementary Table 7: Model output of the association between GABA/Cr and Glu/Cr with covariates

| ROI | Variable | β | t | n participants | P (uncorrected) | P (Bonferroni corrected) |
|---|---|---|---|---|---|---|
| L DLPFC | **Glu/Cr** | 0.40 | 4.25 | 101 | $4.99x10^{-5}$ | $2.99x10^{-4}$ |
| | $Age^{-1}$ | -0.18 | -1.94 | 101 | 0.055 | 0.33 |
| | Frac GM | 0.30 | 3.32 | 101 | 0.0013 | 0.0078 |
| | Sex | -0.21 | -1.17 | 101 | 0.24 | 1.00 |
| R DLPFC | **Glu/Cr** | 0.37 | 3.84 | 99 | $2.20x10^{-4}$ | 0.0013 |
| | $Age^{-1}$ | -0.19 | -2.02 | 99 | 0.047 | 0.28 |
| | Frac GM | -0.12 | -1.20 | 99 | 0.23 | 1.00 |
| | Sex | -0.22 | -1.15 | 99 | 0.26 | 1.00 |
| ACC | **Glu/Cr** | 0.47 | 5.54 | 122 | $1.88x10^{-7}$ | $1.13x10^{-6}$ |
| | $Age^{-1}$ | 0.10 | 1.21 | 122 | 0.23 | 1.00 |
| | Frac GM | -0.0037 | -0.047 | 122 | 0.96 | 1.00 |
| | Sex | 0.21 | 1.35 | 122 | 0.18 | 1.00 |
| MPFC | **Glu/Cr** | 0.47 | 5.47 | 112 | $2.98x10^{-7}$ | $1.79x10^{-6}$ |
| | $Age^{-1}$ | 0.022 | 0.26 | 112 | 0.80 | 1.00 |
| | Frac GM | 0.055 | 0.64 | 112 | 0.53 | 1.00 |
| | Sex | 0.031 | 0.19 | 112 | 0.85 | 1.00 |
| L AIns | **Glu/Cr** | 0.53 | 5.69 | 115 | $1.06x10^{-7}$ | $6.36x10^{-7}$ |
| | $Age^{-1}$ | -0.04 | -0.49 | 115 | 0.63 | 1.00 |
| | Frac GM | -0.11 | -1.37 | 115 | 0.18 | 1.00 |
| | Sex | -0.11 | -0.65 | 115 | 0.52 | 1.00 |
| R AIns | **Glu/Cr** | 0.29 | 3.43 | 123 | $8.38x10^{-4}$ | 0.0050 |
| | **$Age^{-1}$** | 0.33 | 3.93 | 123 | $1.46x10^{-4}$ | $8.76x10^{-4}$ |
| | Frac GM | -0.07 | -0.84 | 123 | 0.40 | 1.00 |
| | Sex | -0.30 | -1.91 | 123 | 0.06 | 0.36 |

Supplementary Table 8: Glu/Cr and GABA/Cr data quality by age

| ROI | Variable | β | t | p |
|---|---|---|---|---|
| L & R DLPFC | Glu/Cr CRLB | 0.08 | 1.26 | 0.20 |
| | GABA/Cr CRLB | 0.09 | 1.35 | 0.17 |
| ACC | Glu/Cr CRLB | -0.08 | -0.90 | 0.37 |
| | GABA/Cr CRLB | 0.10 | 1.24 | 0.22 |
| MPFC | Glu/Cr CRLB | 0.09 | 1.05 | 0.30 |
| | **GABA/Cr CRLB** | 0.27 | 3.08 | 0.003 |
| L & R AIns | Glu/Cr CRLB | -0.08 | -1.23 | 0.22 |
| | GABA/Cr CRLB | 0.06 | 0.72 | 0.47 |

Supplementary Table 9: Glu/Cr model comparison

| **ROI** | **Linear Age AIC** | **Inverse Age AIC** | **Quadratic Age AIC** |
|---|---|---|---|
| L & R DLPFC | -75.16 | **-87.94** | -61.07 |
| ACC | -110.87 | -109.37 | -109.06 |
| MPFC | -49.84 | -49.51 | -47.97 |
| L & R AIns | -301.76 | **-314.80** | -287.21 |

Supplementary Table 10: GABA/Cr model comparison

| **ROI** | **Linear Age AIC** | **Inverse Age AIC** | **Quadratic Age AIC** |
|---|---|---|---|
| L & R DLPFC | -295.31 | **-307.04** | -279.42 |
| ACC | -297.88 | -298.54 | -296.44 |
| MPFC | -225.91 | -226.34 | -224.40 |
| L & R AIns | -549.58 | **-563.54** | -534.52 |

Supplementary Table 11: Glutamate/GABA ratio model comparison

| ROI | Linear Age AIC | Inverse Age AIC | Quadratic Age AIC |
|---|---|---|---|
| L & R DLPFC | 420.03 | **407.98** | 431.91 |
| ACC | 180.30 | **179.98** | 182.13 |
| MPFC | 190.52 | 190.48 | 192.49 |
| L & R AIns | 454.09 | **442.51** | 467.01 |
